# Sympathetic Vasomotion Reflects Catheter-based Radiofrequency Renal Denervation

**DOI:** 10.1101/2025.03.17.643831

**Authors:** Peter Ricci Pellegrino, Irving H. Zucker, Yiannis S. Chatzizisis, Han-Jun Wang, Alicia M. Schiller

**Affiliations:** Department of Anesthesiology, University of Nebraska Medical Center, Omaha, NE 68198; Department of Cellular and Integrative Physiology, University of Nebraska Medical Center, Omaha, NE 68198; Division of Cardiovascular Medicine, Department of Medicine, University of Miami, Miami, FL 33101; College of Medicine, University of Nebraska Medical Center, Omaha, NE 68198

## Abstract

The field of renal denervation remains challenged by the inability to confirm successful ablation of the targeted renal sympathetic nerves. The availability of technology to measure regional blood flow in real time makes sympathetic control of the renal vasculature a logical endpoint to assess effective renal denervation, but autoregulatory mechanisms mask effects on mean renal blood flow. We hypothesized that renal sympathetic vasomotion, a novel marker of rhythmic sympathetic control, reflects successive rounds of catheter-based radiofrequency renal denervation. To test this, ten pigs underwent unilateral surgical renal denervation, recovered for at least seven days, and then underwent four successive rounds of catheter-based radiofrequency denervation of the contralateral kidney. Bilateral renal blood flow velocity and abdominal aortic pressure were measured before and after ablations to assess renal vasomotion. Prior to catheter-based denervation, the renal vasomotion profiles of the innervated and surgically denervated kidneys differed significantly (P < 0.005). Ablation of the largest renal branch artery reduced renal sympathetic vasomotion by 52%. Ablation of the remaining renal branch arteries reduced sympathetic vasomotion 95% from baseline and eliminated the statistical differences between surgically and catheter denervated kidneys. Two additional rounds of catheter denervation of the main renal artery did not consistently decrease renal sympathetic vasomotion magnitude any further. These results indicate that renal sympathetic vasomotion could provide intraprocedural feedback for interventionalists performing catheter-based renal denervation and thereby improve the efficacy, safety, and consistency of this antihypertensive intervention.

## Introduction

The sympathetic nervous system is a master regulator of body homeostasis, and maladaptive sympathetic activation is a driver of the pathophysiology of hypertension^1^, heart failure^2^, and chronic kidney disease^3^. Despite the importance of the sympathetic nervous system in health and disease, clinicians lack a method for assessing regional sympathetic outflow, negatively affecting therapies targeting the sympathetic nervous system and management of chronic sympathoexcitatory diseases. We previously described a novel method leveraging simultaneous recordings of blood pressure and flow to quantify rhythmic sympathetic vascular control and thereby sympathetic outflow, termed sympathetic vasomotion, that could potentially address this clinical gap^4^.

Sympathetic vasomotion has immediate implications for the developing field of renal denervation. Decades of preclinical work established the importance of the renal sympathetic nerves in blood pressure regulation^5^, and the development of clinical intravascular neuroablation devices translated this knowledge into an exciting interventional treatment for hypertension. Despite promising early-stage studies^6,7^, the first pivotal clinical trial failed to meet its primary efficacy endpoint, prompting improvements in device and trial design^8^. Even with these advances, the most recent clinical trial results are mixed. In trials where antihypertensive pharmacotherapy is rigidly controlled or excluded, catheter-based renal denervation lowers blood pressure^9–13^. In the presence of standard-of-care medical therapy, the added value of renal denervation is less clear^14–18^. These nuanced clinical trial results highlight the need for technologies like sympathetic vasomotion that can be used to select treatment responders, validate interventional success, and augment confidence in the technology for patients, practitioners, and other stakeholders.

Our previous study demonstrated that both surgical renal denervation in rabbits and neuraxial sympathetic blockade in swine decrease renal sympathetic vasomotion, but it did not use clinically relevant renal denervation devices or establish a crucial dose-response relationship between renal denervation and renal sympathetic vasomotion^4^. This study was designed to address these shortcomings by performing a regimented catheter-based denervation of pigs that had previously undergone surgical denervation of the contralateral kidney. We hypothesized that successive rounds of catheter-based radiofrequency renal denervation would decrease renal sympathetic vasomotion in a dose-dependent manner.

## Methods

All methods are further detailed in the Online Supplement.

### Data and Materials Availability

The data and source code from this study are available from the corresponding author upon reasonable request.

### Unilateral Surgical Renal Denervation

Experiments were carried out on ten male domestic swine weighing between 44 and 59 kg. Pigs were induced with intramuscular administration of tiletamine, zolazepam, and atropine, endotracheally intubated, and anesthetized with inhaled isoflurane for the duration of the surgery. After shaving and preparing the operative side, the surgical site was anesthetized with 0.5% bupivacaine, and a flank incision was made. The operative kidney was exposed via a retroperitoneal approach, and the renal artery and vein were carefully dissected using glass rods. All neural tissue was stripped off the vessels with microscopic guidance, and the vessels were painted with 10% phenol. The surgical site was then closed in layers. Post-operative pain was treated with scheduled intramuscular buprenorphine and carprofen.

### Contralateral Catheter-based Denervation with Bilateral Vasomotion Monitoring

Pigs were allowed to recover at least seven days from unilateral renal denervation surgery, after which they underwent catheter-based radiofrequency renal denervation of the contralateral kidney. Anesthesia was induced with intramuscular tiletamine and zolazepam and maintained with a ketamine and midazolam infusion after endotracheal intubation. The pigs were left spontaneously ventilating with continuous end-tidal capnography, pulse oximetry, three-lead electrocardiogram, and invasive arterial blood pressure monitoring. Bilateral femoral arterial access was established percutaneously, and pulsed-wave Doppler flow wires were advanced into each renal artery (Figure 1, Figure S1). Heparin was administered and activated clotting time maintained above 250 seconds for thromboprophylaxis for the duration of the experiment. Measurements of abdominal aortic pressure and bilateral renal blood flow velocity were obtained for offline sympathetic vasomotion analysis. The kidney with intact innervation then underwent four successive rounds of catheter-based radiofrequency ablation (Symplicity Spyral, Medtronic plc) in the following systematic, distal-to-proximal fashion: first, the largest branch artery, second, any remaining branch arteries, third, the distal main renal artery, last, the proximal main renal artery (Figure 1, Figure S2). Each round of ablation was followed by measurement of aortic pressure and bilateral renal blood flow velocity. After four rounds of catheter-based renal denervation, the anesthetic infusion was discontinued, and the pigs emerged from anesthesia.

**Fig. 1.**
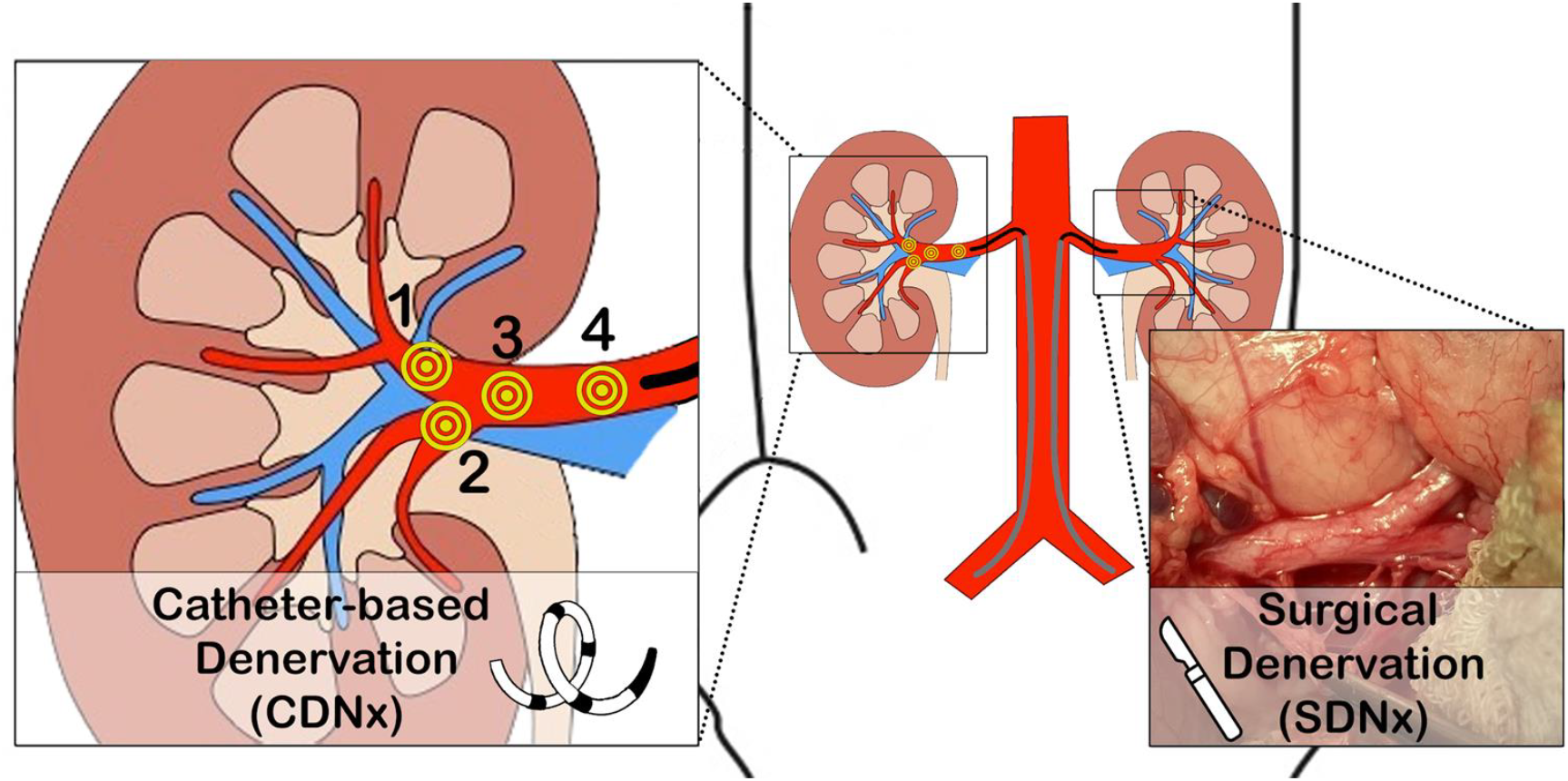
Experimental Paradigm. Pigs underwent unilateral surgical renal denervation (SDNx) two weeks prior to systematic catheter-based denervation (CDNx) of the contralateral kidney. Bilateral renal flow velocity and abdominal aortic pressure were measured simultaneously to compute measures of sympathetic vasomotion.

### Renal Norepinephrine Measurements

Two to four days after catheter-based denervation, pigs were again anesthetized with intramuscular tiletamine and zolazepam, endotracheally intubated, deeply anesthetized with isoflurane 4%, and then euthanized. Both kidneys were collected, snap frozen, and stored at −80 degrees Celsius. For comparison, fully innervated (INV) kidneys from eight swine from both pilot studies and a prior cohort of pigs were collected in the same manner. These samples were then shipped on dry ice to the laboratory of Gregory Fink, Ph.D., which specializes in high-performance liquid chromatography for the measurement of tissue norepinephrine. This was performed in quintuplicate on homogenized renal tissue as described previously^19^. The median norepinephrine concentration for each kidney expressed as picograms of norepinephrine per gram of renal tissue was used for analysis.

### Data Analysis

All processing was done offline; renal sympathetic vasomotion was not calculated in real time to avoid any potential of biasing the procedural team. The relationship between arterial pressure (AP) and the resistive component of renal blood flow velocity (RBFV) was characterized using time-frequency analysis, yielding vasomotion profile occurrence histograms, in a method introduced previously^20^. The three components of the AP-RBFV relationship have distinct physiologic underpinnings. Admittance gain represents active modulation of vascular resistance at a given frequency. Phase shift quantifies differences in the timing of oscillations in blood pressure and blood flow; negative phase shift behavior (i.e., arterial pressure oscillations precede the modulation of renal vascular resistance) baroreflex-mediated vascular control. Coherence, which was averaged over three independent wavelet lengths, represents how closely flow velocity passively follows pressure at a given time and frequency and is thus inversely related to total active vasomotion. Total renal vasomotion difference, a composite measure of the differences from all transfer function components, was calculated as the square root of the sum of squares for the differences in admittance gain, phase shift, and coherence. Renal sympathetic vasomotion magnitude was derived from individual occurrence histograms selecting vasomotion behavior identified as innervated or denervated via cluster-mass based statistical testing on vasomotion data prior to catheter-based denervation.

### Statistical Analysis

Statistical testing of simple, normally distributed measures (e.g., arterial pressure, heart rate, renal blood flow velocity) was conducted as a repeated measures ANOVA with appropriate within-subjects factors (e.g., ablation number, denervation modality) and α = 0.05. Such data are displayed as mean ± standard deviation. Statistical testing of renal norepinephrine data was performed with Wilcoxon signed-rank test for surgical (SDNx) versus catheter-based denervation (CDNx) and Mann-Whitney U test for INV versus denervated kidneys; a Holm-Sidak correction was used to correct for multiple comparisons. Group vasomotion profile occurrence histograms are displayed as mean data for each group. Vasomotion difference maps computed as t-statistics for each independent variable pair (e.g., frequency-admittance gain bin) using two-tailed, paired t-test are displayed to convey directionality, magnitude, and consistency of differences but not statistical significance *per se*. Due to the high dimensionality of the vasomotion occurrence data, statistical testing was performed with non-parametric cumulative difference mass testing, which allows for *a priori* significance testing of non-independent, multidimensional data while addressing the inherent multiple comparisons problem as described previously^20,21^. The cumulative differences for both the actual data and the surrogate shuffled data are displayed along with P values.

## Results

### Denervation Validation

Each pig underwent four rounds of successive renal radiofrequency neurotomy, with an average of 12.8 total sites ablated (Table S1). Both surgical and catheter-based renal denervation resulted in significant decreases in renal norepinephrine content; there was no significant difference in norepinephrine content between surgically and catheter-based denervation (Figure 2).

**Fig. 2.**
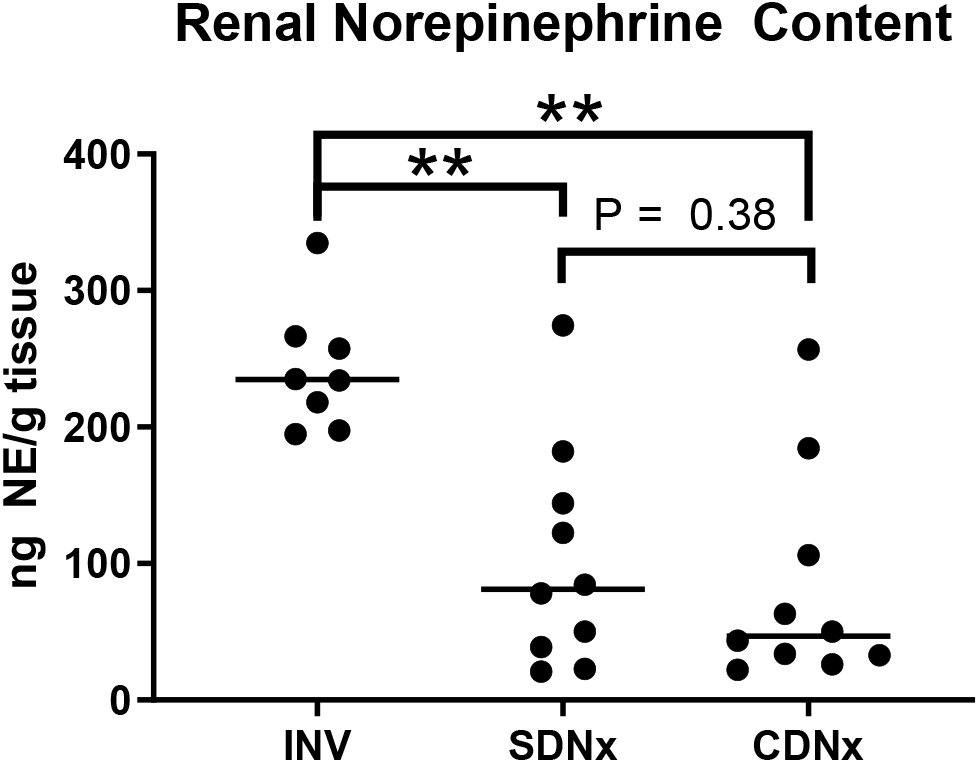
Renal Norepinephrine Content. Both renal denervation modalities significantly decreased renal norepinephrine content relative to kidneys from pigs with intact renal innervation. Lines depict median norepinephrine concentration for each group.

### Hemodynamics

Mean arterial pressure and heart rate did not significantly change over the duration of the experiment (Figure S3A and S3B). Bilateral renal blood flow velocity was similar between the surgically denervated kidneys and the kidneys undergoing catheter-based denervation with no statistically significant effects of denervation modality, ablation number, or the interaction between these factors (Figure S3C).

### Admittance Gain

Admittance gain is the vasomotion component that represents the magnitude of transduction of arterial pressure oscillations into oscillations in blood flow velocity. With respect to admittance gain profiles, the surgically denervated kidney and the contralateral kidney differed significantly (P < 0.005) prior to catheter-based denervation. Largest branch denervation diminished this difference both in terms of magnitude and statistical significance, and the difference was eliminated after complete branch denervation (2 rounds of ablation). Occurrence histograms showed more high admittance gain behavior in the innervated kidney prior to catheter-based denervation that was eliminated after complete branch artery denervation (Figure 3B-F, Figure S4). Subsequent rounds of radiofrequency ablation of the distal and proximal main renal artery did not significantly impact admittance gain.

**Fig. 3.**
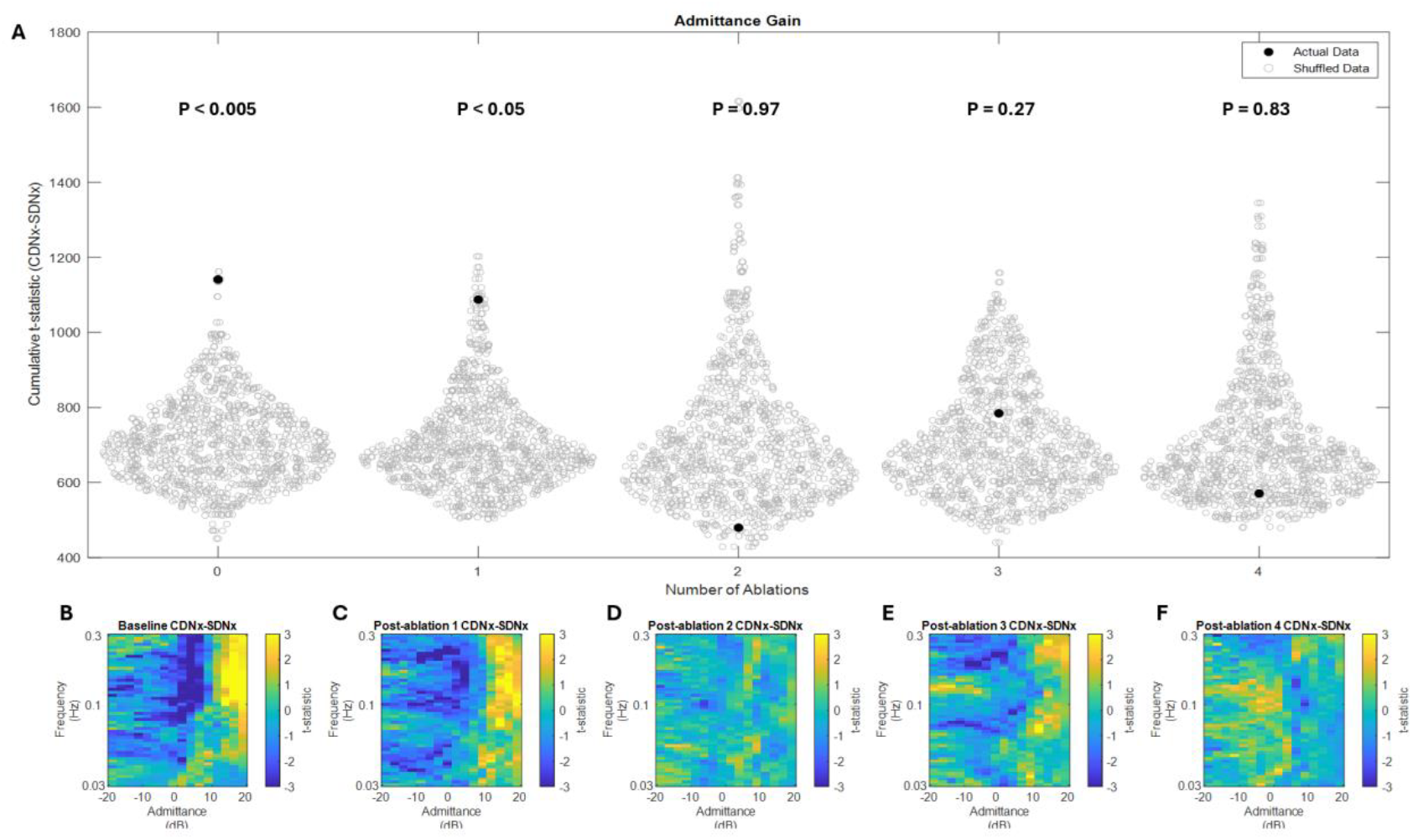
Admittance Gain Differences. (A) Non-parametric mass-based statistical analysis shows significant differences between innervated and surgically denervated kidneys in admittance gain behavior that are eliminated by catheter-based renal denervation. (B) The occurrence difference profile in the baseline state shows high admittance gain behavior in the innervated kidney relative to the surgically denervated kidney. (C) This high admittance gain behavior is abrogated by largest branch artery denervation and (D) eliminated by complete branch artery denervation with no further changes with (E) distal main artery ablation and (F) proximal main artery ablation.

### Phase Shift

Phase shift quantifies the temporal relationship between oscillations in blood pressure and blood flow and provides insight into the directionality of control mechanisms, with negative phase shift behavior indicative of baroreflex-mediated vascular control. The phase shift component of the vasomotion profiles also differed significantly prior to catheter-based denervation (Figure 3A). This difference was no longer statistically significant after largest branch radiofrequency ablation although a trend remained (P = 0.08); this trend was eliminated by subsequent denervation of the remaining branch arteries. Occurrence histograms showed more baroreflex-mediated, negative phase shift behavior in the innervated kidney prior to denervation with reduction in this phenomenon after largest branch denervation and elimination of this difference after catheter-based denervation of the remaining branch arteries (Figure 4B-F, Figure S5). Catheter-based denervation of the distal and proximal main artery did not further impact the phase shift component of vasomotion after full branch denervation.

**Fig. 4.**
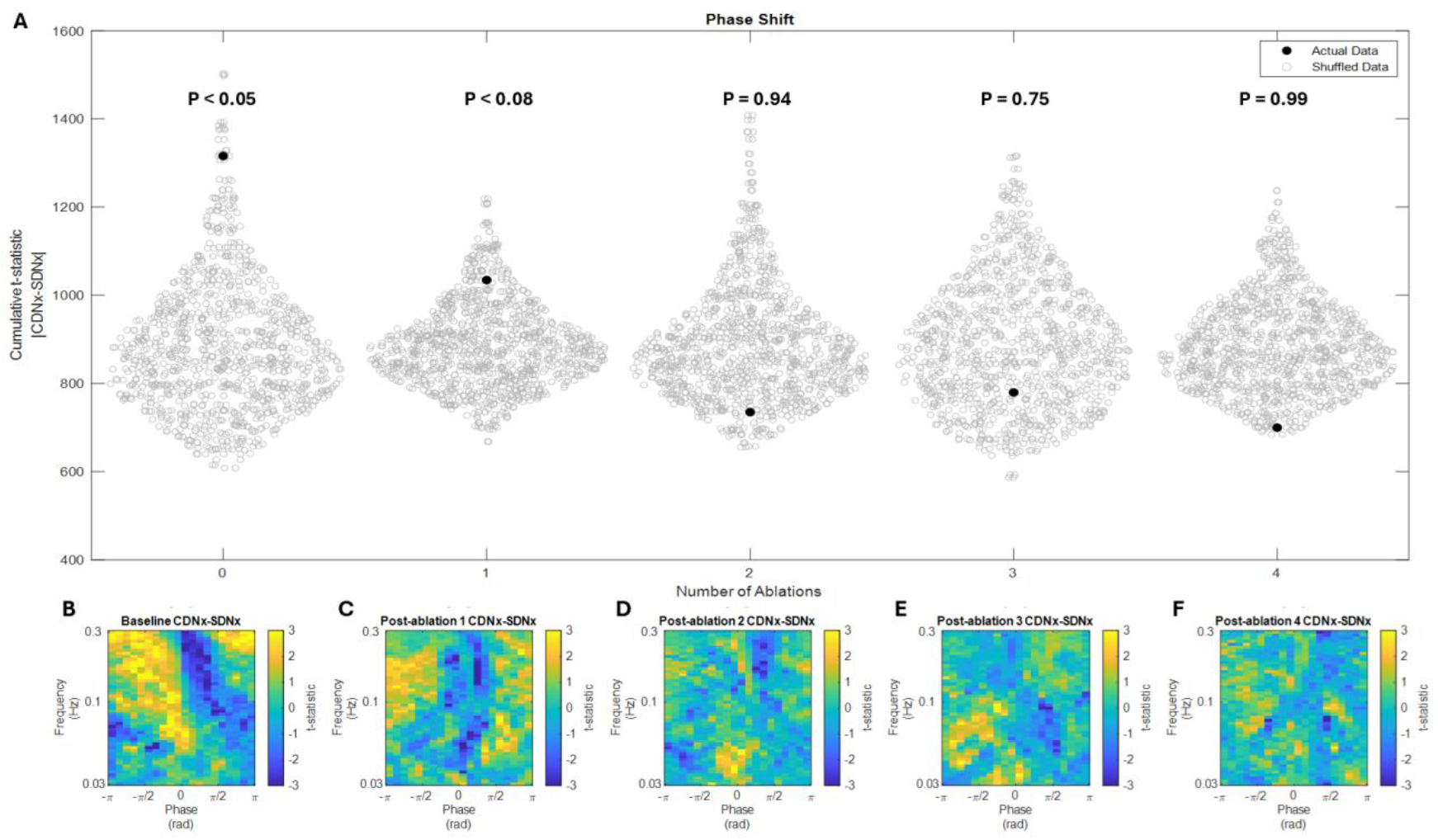
Phase Shift Differences. (A) Non-parametric mass-based statistical analysis shows significant differences between innervated and surgically denervated kidneys in phase shift behavior that are eliminated by catheter-based renal denervation. (B) The occurrence difference profile in the baseline state shows more negative phase shift vasomotion in the innervated kidney relative to the surgically denervated kidney, consistent with baroreflex control. (C) This negative phase shift is abrogated by largest branch artery denervation and (D) eliminated by complete branch artery denervation with no further changes with (E) distal main artery ablation and (F) proximal main artery ablation.

### Coherence

Coherence quantifies the consistency in the pressure-flow relationship and correlates inversely with active vascular modulation. Coherence differed significantly between the surgically denervated and the fully innervated kidney prior to catheter-based denervation (Figure 5A). The magnitude and non-parametric significance of this difference decreased after radiofrequency ablation of the largest branch artery, falling below statistical significance (P = 0.26), and decreased further with subsequent denervation of the remaining branch arteries. Prior to catheter-based denervation, the surgically denervated kidney exhibited more high coherence behavior, consistent with the removal of an active vascular control mechachnism by surgical denervation (Figure 5B). This difference was eliminated by subsequent catheter-based radiofrequency ablation.

**Fig. 5.**
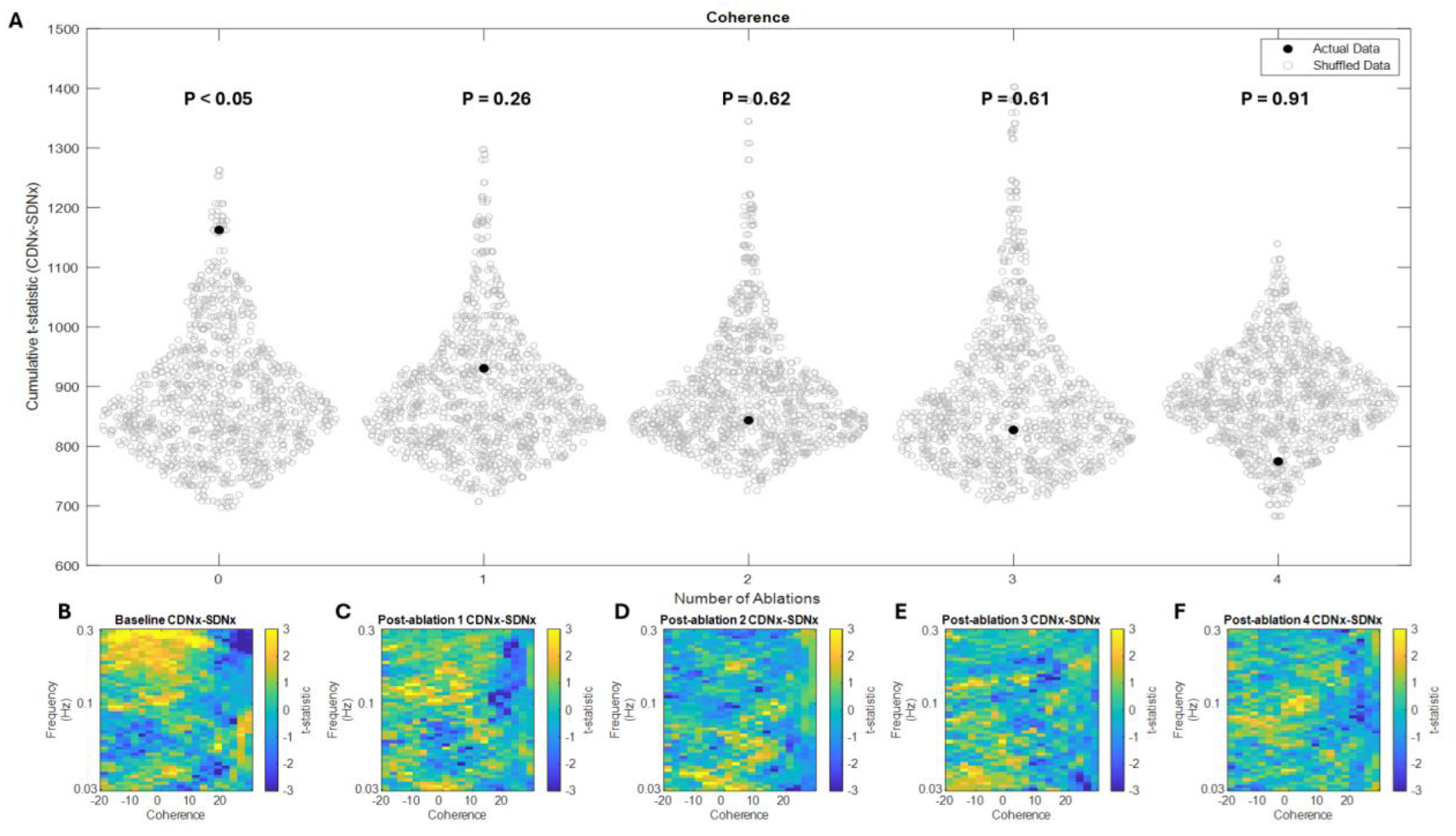
Coherence Differences. (A) Non-parametric mass-based statistical analysis shows significant differences between innervated and surgically denervated kidneys in coherence behavior that are eliminated by catheter-based renal denervation. (B) The occurrence difference profile in the baseline state shows more low coherence vasomotion in the innervated kidney relative to the surgically denervated kidney, consistent with active vascular control. (C) This difference is statistically eliminated by largest branch artery denervation with a subtle further decrease with complete branch artery denervation. (E) Distal main artery ablation and (F) proximal main artery ablation do not further reduce differences in coherence behavior between the catheter-based and surgically denervated kidneys.

### Total Vasomotion and Renal Sympathetic Vasomotion Magnitude

Two measures were used to aggregate the vasomotion data from the three components of the time-varying transfer function, total vasomotion profile difference and renal sympathetic vasomotion magnitude. The total vasomotion profile difference is a statistical group measure that reflects the statistical differences across all components of the time-varying pressure-resistive flow transfer function. Prior to catheter-based denervation, total vasomotion differed significantly between the surgically denervated kidney and the contralateral innervated kidney (Figure 6A). This difference was decreased by radiofrequency denervation of the largest branch artery and eliminated by complete branch denervation with no obvious benefit from additional rounds of radiofrequency ablation in the main renal artery. Renal sympathetic vasomotion magnitude is an individual measure of sympathetic vascular control that reflects how this technology could be used clinically on a patient-by-patient basis. As expected, renal sympathetic vasomotion is stable in the SDNx kidneys while the contralateral kidney undergoes catheter-based denervation (Figure 6B). Radiofrequency ablation of the largest renal branch artery results in a 52% decrease in renal sympathetic vasomotion magnitude, and ablation of the remaining branch arteries reduces sympathetic vasomotion magnitude an additional 43% (95% from baseline). Subsequent radiofrequency ablations of the distal and proximal main renal artery do not consistently decrease renal sympathetic vasomotion magnitude any further. The interaction between denervation modality and ablation number statistically corroborates the specificity of the effect of renal denervation on sympathetic vasomotion of CDNx kidneys.

**Fig. 6.**
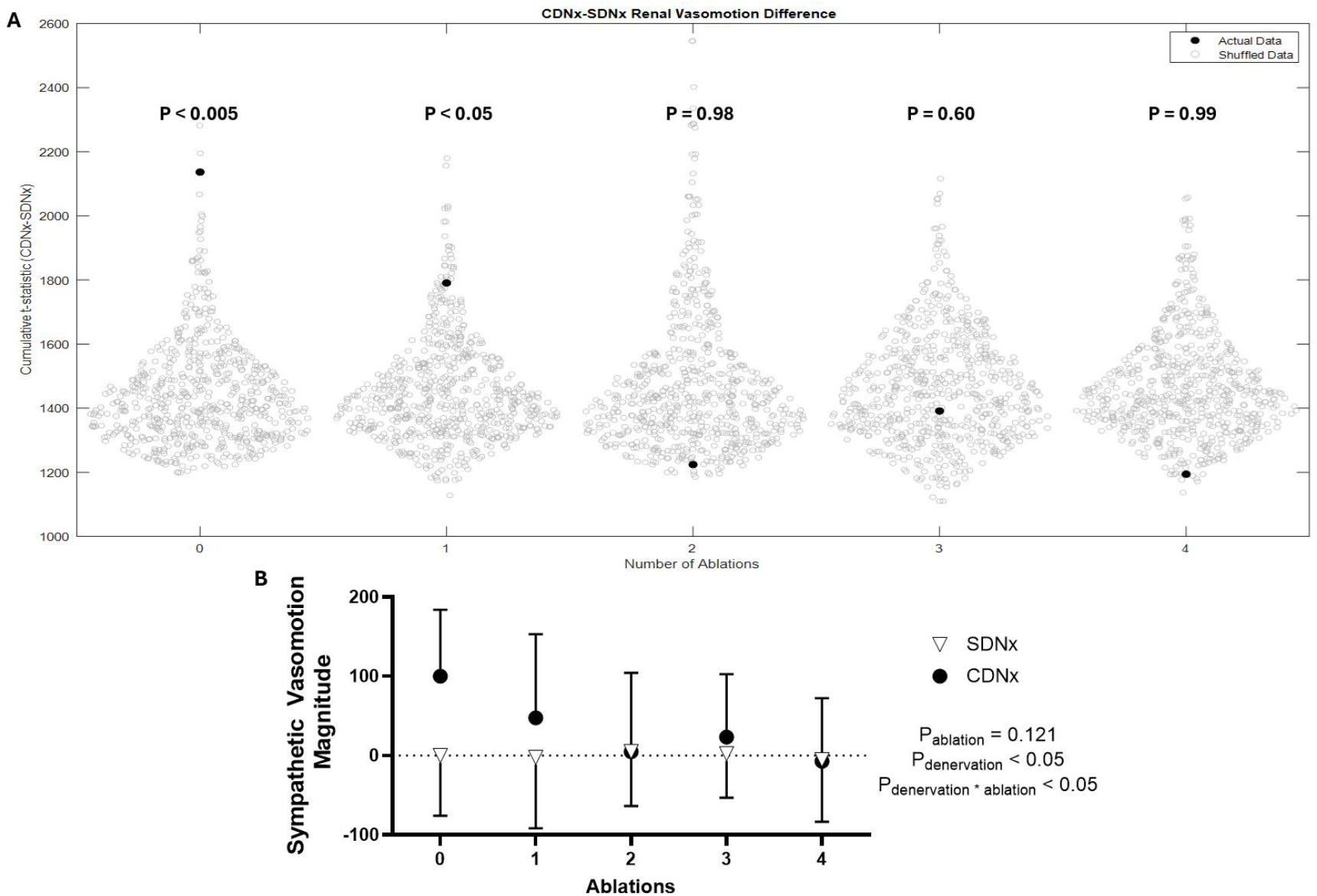
Aggregate measures of renal sympathetic vasomotion. (A) Non-parametric mass-based statistical analysis shows significant differences between innervated and surgically denervated kidneys in total vasomotion behavior that are decreased with catheter-based radiofrequency of the largest branch artery and eliminated by complete branch artery ablation. Further ablation of the distal and proximal main renal artery does not affect cumulative vasomotion differences between the kidneys. (B) Catheter-based denervation of the renal branch arteries decreases renal sympathetic vasomotion in CDNx kidneys with no further benefit from main artery lesioning. This measure remains stable in SDNx kidneys.

## Discussion

These data show that renal sympathetic vasomotion reflects the renal denervation caused by successive rounds of catheter-based radiofrequency ablation. Moreover, the sympathetic vasomotion findings are corroborated by tissue norepinephrine levels; both show that catheter-based radiofrequency ablation reduces renal sympathetic innervation as much as surgical renal denervation. This serves as a proof-of-concept for the use of renal sympathetic vasomotion as an intraprocedural monitoring tool for catheter-based renal denervation.

### Real-time Intraprocedural Feedback for Renal Denervation

The need for intraprocedural feedback is well-recognized by interventionists, and thus several other approaches have been deployed for this purpose. Renal norepinephrine spillover was used in an earlier proof-of-concept clinical trial to validate the first-generation radiofrequency denervation device^7^ and showed the heterogeneity of renal denervation with this device^23^. Unfortunately, this technique is time-consuming, invasive, and unable to be implemented on a widespread clinical basis. Another studied approach is the use of electrical stimulation to assess the afferent-mediated pressor response or the efferent-mediated renovascular constriction before and after renal denervation. The proponents of this approach assert that it allows for the differentiation between pressor and depressor loci, potentially increasing the antihypertensive effect of renal denervation or decreasing the number of lesions^24,25^. Mapping introduces a significant degree of procedural complexity, and, since stimulation of the renal afferents is painful, requires deep sedation or general anesthesia. The ability to passively assess renal denervation efficacy is an advantage of renal sympathetic vasomotion as an intraprocedural endpoint for renal denervation.

This much-needed feedback for the procedure could be used on a patient-by-patient basis (e.g., patients with anomalous anatomy) or leveraged to compare different denervation strategies in humans. For clinicians performing renal denervation procedures, this study highlights the importance of branch denervation and raises questions about the marginal benefit of main artery denervation after complete branch artery neurotomy, at least in swine, and this finding is consistent with tissue norepinephrine data from prior swine studies and recent clinical studies^26^. Whether a streamlined procedure could offer patients similar benefit with less procedural time and risk could be addressed with clinical studies enhanced by sympathetic vasomotion monitoring. While calculation of renal sympathetic vasomotion was performed offline in order to avoid any potential biasing of the procedural team, the processing is not computationally intensive and can easily be done in real time to provide the immediate feedback needed for intraprocedural decision-making.

### Implications Beyond Intraprocedural Monitoring for Renal Denervation

In addition to intraprocedural monitoring, measurement of renal sympathetic vasomotion could be very important for preprocedural patient selection and long-term monitoring for reinnervation. Essential hypertension is a heterogenous disorder, and the variability in the blood pressure response to renal denervation among patients has been a point of criticism. For other percutaneous neurotomy procedures, like medial branch radiofrequency ablation for axial spine pain, the standard-of-care is to perform diagnostic steps prior to therapeutic procedures to improve patient outcomes^27^. Preprocedural measurement of renal sympathetic vasomotion may be useful for patient selection, serving as a biomarker for those with high renal sympathetic tone who would be expected to respond more substantially to renal denervation. This could be done using non-invasive measures of renal blood flow (e.g., transabdominal ultrasound) and blood pressure. Moreover, many animal studies suggest that renal reinnervation occurs even after surgical denervation^28,29^, and renal sympathetic vasomotion monitoring could be a useful method of identifying patients who have evidence of functional reinnervation and thus might benefit from repeat ablation. This technology may also allow for more targeted use of renal denervation beyond hypertension. For example, our recent work suggests that elevated renal sympathetic outflow drives the progression of cardiorenal type 2 syndrome after myocardial infarction in rats.^30^ Renal sympathetic vasomotion could be used to identify patients with elevated renal sympathetic tone after myocardial infarction, so they could undergo renal denervation to preserve renal function. Finally, the ability to quantify sympathetic outflow to a given vascular bed has implications beyond the field of renal denervation, including diagnosis and treatment of other diseases with sympathetic dysregulation including heart failure, chronic kidney disease, dysautonomia, complex regional pain syndrome, and shock states

### Limitations and Strengths of the Study

Our study has several limitations. We were unable to identify a reliable surrogate for functional renal innervation despite testing multiple interventions in pilot studies, including the renovascular response to femoral arterial adenosine, the hemodynamic response to renal arterial capsaicin, percutaneous electrical stimulation of sympathetic paravertebral ganglia, and hypoxia. The challenges with using these reflexes to assess innervation status further highlights the importance of a passive measure like renal sympathetic vasomotion. Furthermore, there was substantial variability in the effect of both catheter-based and surgical renal denervation on tissue norepinephrine levels. Some of this may have arisen due to circulating norepinephrine as these animals were not perfused prior to euthanasia. The median decrease in tissue norepinephrine levels was similar to those in other studies^22^. Some potential issues with the inherent variability in denervation magnitude could be addressed by future studies in which the effects of successive rounds of catheter-based denervation on renal sympathetic vasomotion are compared to a fully innervated kidney, providing a logical contrapositive to this study. The strengths of the study include the use of a clinically relevant renal denervation device and clinical intravascular pressure-flow monitoring devices. Additionally, the utilization of the contralateral kidney as a control eliminates confounding, time-dependent effects like those of anesthetic depth and changes in neurohumoral activity.

### Conclusions

In summary, sympathetic vasomotion, a novel measure of sympathetic vascular control, demonstrates a dose-response relationship with catheter-based renal denervation in swine. This highlights its potential utility as an intraprocedural feedback measure for interventionalists performing this procedure. The translation of this work to patients could improve the efficacy, safety, and consistency of renal denervation and has implications for other diseases in which sympathetic dysregulation plays a causal role.

## Acknowledgments

The authors would like to thank John Lof, Elizabeth Stolze, Kaye Talbitzer, Hannah Garver, and Gregory Fink, Ph.D., for their technical assistance.

